# The balance of mitochondrial fission and fusion in cortical axons depends on the kinases SadA and SadB

**DOI:** 10.1101/2021.07.18.452621

**Authors:** Danila Di Meo, Priyadarshini Ravindran, Pratibha Dhumale, Andreas W. Püschel

## Abstract

Neurons are highly polarized cells that display characteristic differences in the organization of their organelles in axons and dendrites. Mitochondria are of particular importance for neuronal homeostasis due to their high metabolic demand. The kinases SadA and SadB (SadA/B) promote the formation of distinct axonal and dendritic extensions during the development of cortical and hippocampal neurons. Here, we show that SadA/B are required for the axon-specific dynamics of mitochondria. The interaction with Ankyrin B (AnkB) stimulates the activity of SadA/B that function as regulators of mitochondrial dynamics through the phosphorylation of Tau. Suppression of SadA/B or AnkB in cortical neurons induces the elongation of mitochondria by disrupting the balance of fission and fusion. The normal dynamics of axonal mitochondria could be restored by mild actin destabilization. Thus, the elongation after a loss of SadA/B results from an excessive stabilization of actin filaments and reduction of Drp1 recruitment to mitochondria.

## Introduction

Neurons are highly polarized cells with morphologically and molecularly distinct axonal and somato-dendritic compartments that display characteristic differences in the organization of their organelles (Britt et al., 2016; Schelski and Bradke, 2017). Of particular importance for neuronal homeostasis are mitochondria. Defects in their morphology or dynamics lead to neurodevelopmental diseases and neurodegeneration (Baum and Gama, 2021). Mitochondria are dynamic organelles that are continuously remodeling their morphology by fission and fusion (Fenton et al., 2021; Moore, 2018). Mitochondrial dynamics is a finely tuned process that regulates their size and distribution and is essential for their function (Lee et al., 2018; Sheng and Cai, 2012). Fusion of the outer mitochondrial membrane (OMM) is mediated by Mitofusins while fission depends on the cytoplasmic GTPase Dynamin-related protein 1 (Drp1) that is recruited to the OMM by receptors like Mitochondrial fission factor (Mff) (Losón et al., 2013). Mitochondria are relatively short and uniform in size in axons while they are more elongated and display lower motility in the somato-dendritic compartment (Cagalinec et al., 2013; Chang et al., 2006; Lewis et al., 2018; Rangaraju et al., 2019; Saxton and Hollenbeck, 2012). Maintaining the size of axonal mitochondria depends on the coupling of fission and fusion and requires Mff (Lewis et al., 2018). Since Mff is present in both axons and dendrites the factors that are responsible for regulating the balance of fission and fusion and the size of mitochondria in the axon remain to be identified.

During the development of the nervous system, axons and dendrites arise from undifferentiated neurites that are extended by newborn unpolarized neurons (Lewis et al., 2013; Namba et al., 2015; Schelski and Bradke, 2017). Upon the establishment of neuronal polarity, one of these neurites is selected to differentiate as the axon, while the remaining neurites acquire the characteristics of dendrites. The kinases SadA and SadB (SadA/B, also known as Brsk2 and Brsk1) function redundantly to promote the formation of distinct axonal and dendritic extensions in cortical and hippocampal neurons (Barnes et al., 2007; Dhumale et al., 2018; Kishi et al., 2005; Müller et al., 2010; Shelly and Poo, 2011). They are also required at later stages for different aspects of neuronal differentiation like the central arborization of sensory axons as well as the maturation of synapses in the central and peripheral nervous system (Barnes et al., 2007; Kishi et al., 2005; Lilley et al., 2013; Shelly and Poo, 2011). They belong to the AMPK family of serine/threonine kinases and are activated by phosphorylation of Thr175 and Thr187, respectively, by the Lkb1/Strad complex (Barnes et al., 2007; Lizcano et al., 2004; Shelly and Poo, 2011; Veleva-Rotse et al., 2014). However, their regulation is cell-type specific. Not all functions of SadA/B depend on Lkb1 and other kinases can also activate them (Bright et al., 2008; Lilley et al., 2013).

A knockout of both *Sada* and *Sadb* severely reduces cortical axon tracts and the size of the cortex, which is not detectable in single knockouts (Dhumale et al., 2018; Kishi et al., 2005). The *Sada*;*Sadb*-deficient cortex shows extensive apoptosis of neurons and a reduction in the number of progenitors. Cultured hippocampal or cortical neurons from *Sada*^-/-^;*Sadb*^-/-^ double knockout embryos fail to form distinct axonal and dendritic processes. Instead, a large proportion extends indeterminate neurites that are positive for both axonal and dendritic markers. The defect in axon formation has been linked to a function in the regulation of microtubule binding proteins (Barnes et al., 2007; Cavallini et al., 2013; Kishi et al., 2005). SadA/B and other AMPK-related kinases like Mark2 or Nuak1 phosphorylate the microtubule-associated protein Tau at Ser262 and Ser356, which attenuates the interaction with microtubules (Biernat et al., 1993; Drewes et al., 1995). However, how Tau regulates microtubule dynamics and contributes to the defects in neuronal polarity after the loss of SadA/B still remains incompletely understood. Tau is generally thought to stabilize microtubules, but recent results indicate that it promotes the elongation of labile microtubule domains without stabilizing them (Chang et al., 2021; Qiang et al., 2018). Tau oligomerizes to form condensations on microtubules that act as barriers for kinesins (Ambadipudi et al., 2017; Chang et al., 2021; Siahaan et al., 2019; Tan et al., 2019; Wegmann et al., 2018). The formation of condensates is promoted by its phosphorylation but does not require it. Tau hyperphosphorylation is a hallmark of Alzheimer’s disease and has been linked to defects in the axonal transport of mitochondria (Chang et al., 2021).

In this study, we identified a novel function of SadA/B kinases as regulators of mitochondrial dynamics that is required for maintaining the typical size of mitochondria in cortical axons. Suppression of SadA/B leads to an elongation of mitochondria by disrupting the balance between fission and fusion. Our results show that the increased length of mitochondria after the loss of SadA/B results from an excessive stabilization of F-actin and an impairment of Drp1 recruitment to the OMM, due to the reduced phosphorylation of Tau. The scaffolding protein Ankyrin B (AnkB) functions upstream of SadA/B to stimulate their phosphorylation.

## Results

### AnkB interacts with SadA and B and stimulates their activation

SadA/B are kept in an auto-inhibited state by the association of their N-terminal kinase domain with the ubiquitin associated (UBA) domain and an auto-inhibitory sequence (AIS) at the C-terminus (Fig. 1A) (Ma et al., 2016; Wu et al., 2015). This auto-inhibition can be released by the interaction with scaffolding proteins (Hung et al., 2007; Ma et al., 2016; Wu et al., 2015). Active SadA and SadB, indicated by phosphorylation at Thr175 (SadA) and Thr186 (SadB), are restricted to the axonal compartment (Barnes et al., 2007; Dhumale et al., 2018). The scaffolding protein AnkB is a component of the submembranous cytoskeleton but also mediates bidirectional motility of organelles by linking them to the dynein/dynactin complex (Galiano et al., 2012; Lorenzo et al., 2014; Qu et al., 2016). To test if AnkB functions as a scaffold for SadA/B we co-expressed them in HEK 293T cells. Immunoprecipitation showed that AnkB interacts with both kinases (Fig. 1B). It binds to the N-terminal fragment containing the kinase and UBA domains but not the C-terminal fragment of SadA and SadB (Fig. 1C).

**Figure 1.**
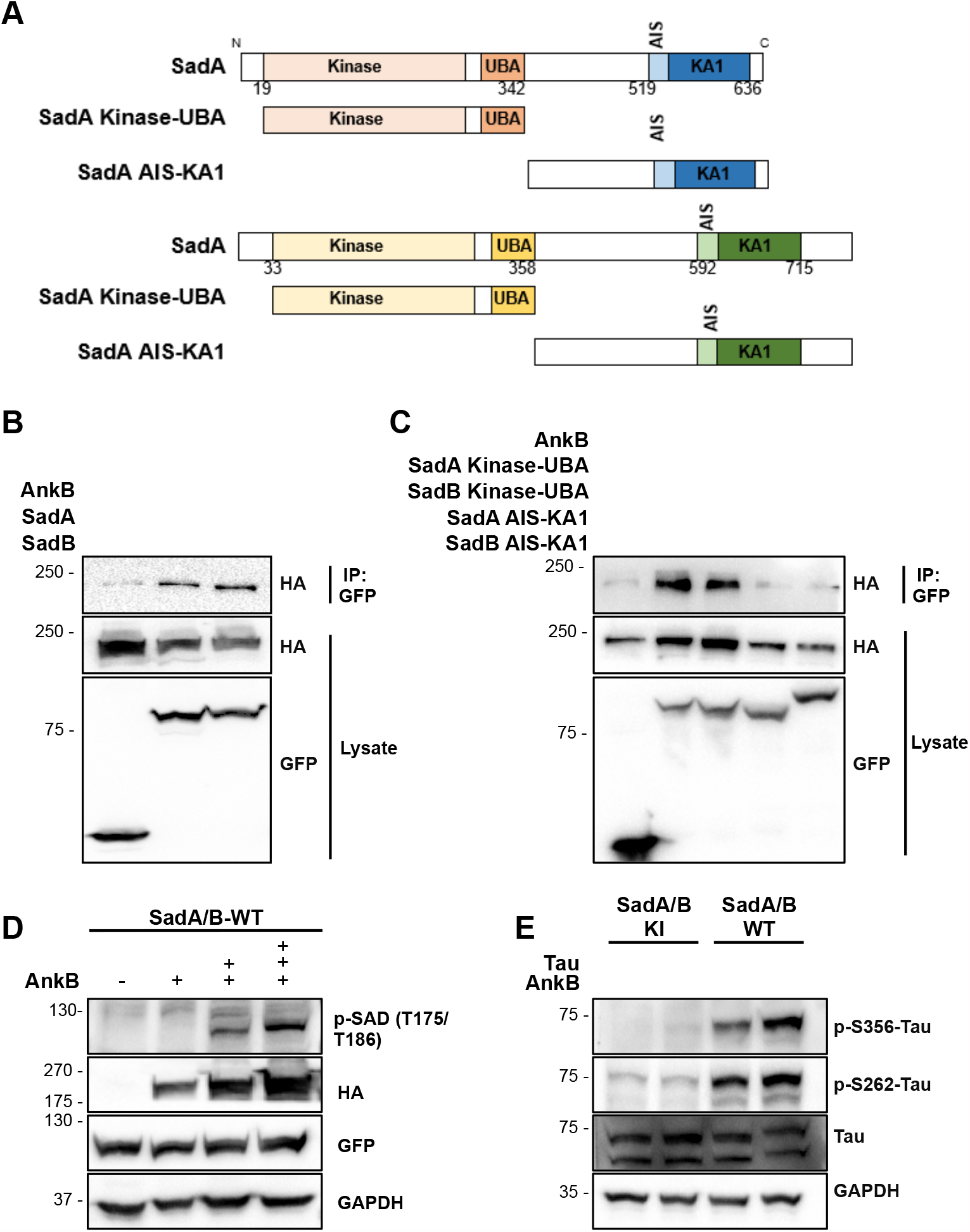
Ankyrin B interacts with SadA/B and stimulates their activation. (A) Schematic representation of SadA and SadB showing the domains used for interaction studies. (B and C) HA-AnkB and GFP-SadA, GFP-SadB (B) or the indicated fragments of SadA and SadB (C) were coexpressed in HEK 293T cells, immunoprecipitated and analyzed by Western blot with the indicated antibodies. (D) Wild type SadA and SadB were coexpressed with increasing amounts of HA-AnkB in HEK 293T cells (SadA/B:AnkB expression vector ratio: 1:5,1:10 and 1:15). SadA/B phosphorylation was analyzed by Western blot with the indicated antibodies. (E) Wild type (SadA/B WT) or inactive SadA and SadB (SadA/B-KI), HA-AnkB and mRuby2-Tau were co-expressed in HEK 293T cells as indicated. Tau phosphorylation was analyzed by Western blot with the indicated antibodies. See also Figures S1.

To investigate whether the interaction with AnkB stimulates the phosphorylation of SadA/B we co-expressed increasing amounts of AnkB with SadA, SadB or non-phosphorylatable SadA-T175A and SadB-T186A in HEK 293T cells. In the absence of AnkB, only low levels of phospho-T175/T186-SadA/B were detectable by Western blot. With increasing expression of AnkB higher levels of phospho-T175/186-SadA/B were observed (Fig. 1D). No signals were detectable for the non-phosphorylatable SadA/B mutants (Fig. S1A). To analyze SadA/B activity we used the microtubule-binding protein Tau as a well-characterized substrate that is phosphorylated at Ser262 and Ser356 (Ercan et al., 2017; Kishi et al., 2005; Yoshida and Goedert, 2012). Wild type SadA/B or inactive SadA/B (SadA/B-KI) were co-expressed with Tau or non-phosphorylatable Tau-S262A, Tau-S356A or Tau-S262/356A in HEK 293T cells and analyzed by Western blot. Only low levels of Tau phosphorylation at Ser262 and Ser356 by endogenous kinases were detectable with SadA/B-KI. Overexpression of wild type SadA/B strongly increased Tau phosphorylation at both sites (Fig. S1B). Notably, co-expression of AnkB stimulates SadA/B activity and increased basal Tau phosphorylation at Ser262 (41±2%) and Ser356 (55±4%) (Fig. 1E, S1C and S1D). Thus, AnkB interacts with SadA/B and promotes their activation possibly by releasing the auto-inhibition to allow the phoshorylation by Lkb1 (Barnes et al., 2007; Lizcano et al., 2004).

### AnkB acts upstream of SadA/B

Neurons from *Sada*^-/-^;*Sadb*^-/-^ double knockout mice fail to form a distinct axon and instead extend several neurites that are relatively uniform in length and positive for both axonal and dendritic markers (Dhumale et al., 2018; Kishi et al., 2005; Müller et al., 2010). To study the function of SadA/B in cortical neurons but avoid the extensive loss of neurons by apoptosis in the *Sada*^-/-^;*Sadb*^-/-^ knockout cortex (Dhumale et al., 2018), we established knockdown vectors for SadA and SadB. The efficiency of the knockdown constructs was confirmed by Western blot after expression in HEK 293T cells (Fig. S2A). Cortical neurons from E18 rat embryos were transfected at 0 d.i.v. with knockdown vectors for SadA and SadB and analyzed at 3 d.i.v. by staining with antibodies for acetylated tubulin as axonal marker and MAP2 as dendritic marker (Fig. S2B). Transfected neurons were marked by expression of GFP from the knockdown vectors. The SadA/B knockdown induced the formation of indeterminate neurites that are positive for both axonal and dendritic markers (control: 13±1%; SadA/B RNAi: 44±1%) as described before (Dhumale et al., 2018; Kishi et al., 2005; Müller et al., 2010). In addition, 33±1% of the neurons extended supernumerary axons that are positive for acetylated tubulin but not MAP2 (Fig. S2B and S2C). This phenotype was rescued by co-expression of RNAi-resistant SadA/B, verifying the specificity of the knockdown constructs (Fig. S2D and S2E). After knockdown of SadA/B, the levels of phospho-S356-Tau detected by Western blot (Fig. S2F) and immunofluorescence were strongly reduced in cortical neurons at 3 d.i.v. (Fig. S2G and S2H).

In order to determine if AnkB is required for axon formation, cortical neurons were transfected with a knockdown vector at 0 d.i.v. The efficiency of the knockdown construct that targets the AnkB death domain shared by the 220 and 440 kDa isoforms (Ayalon et al., 2011) was confirmed by Western blot (Fig. S2I). The knockdown of AnkB increased the number of neurons with indeterminate neurites (control: 6±0.4%, AnkB RNAi: 37±1%) and supernumerary axons (control: 14±1%, AnkB RNAi: 29±1%) similar to the knockdown of SadA/B (Fig. 2A and 2B). Expression of RNAi-resistant AnkB rescued this phenotype confirming the specificity of the knockdown construct (Fig. S2J). The expression of constitutively active SadA (SadAca) and SadB (SadBca) (Barnes et al., 2007; Lizcano et al., 2004; Müller et al., 2010) restored neuronal polarity after the knockdown of AnkB, confirming that it results from a reduced activation of SadA/B (Fig. 2C and 2D; AnkB RNAi: 35±2% indeterminate neurites, 28±1% supernumerary axons; AnkB RNAi+SadA/Bca: 17±2% indeterminate neurites; 17±2% supernumerary axons). Constitutively active SadA/B did not affect neuronal polarity in control neurons. These results indicate that AnkB acts upstream of SadA/B to stimulate their activity.

**Figure 2.**
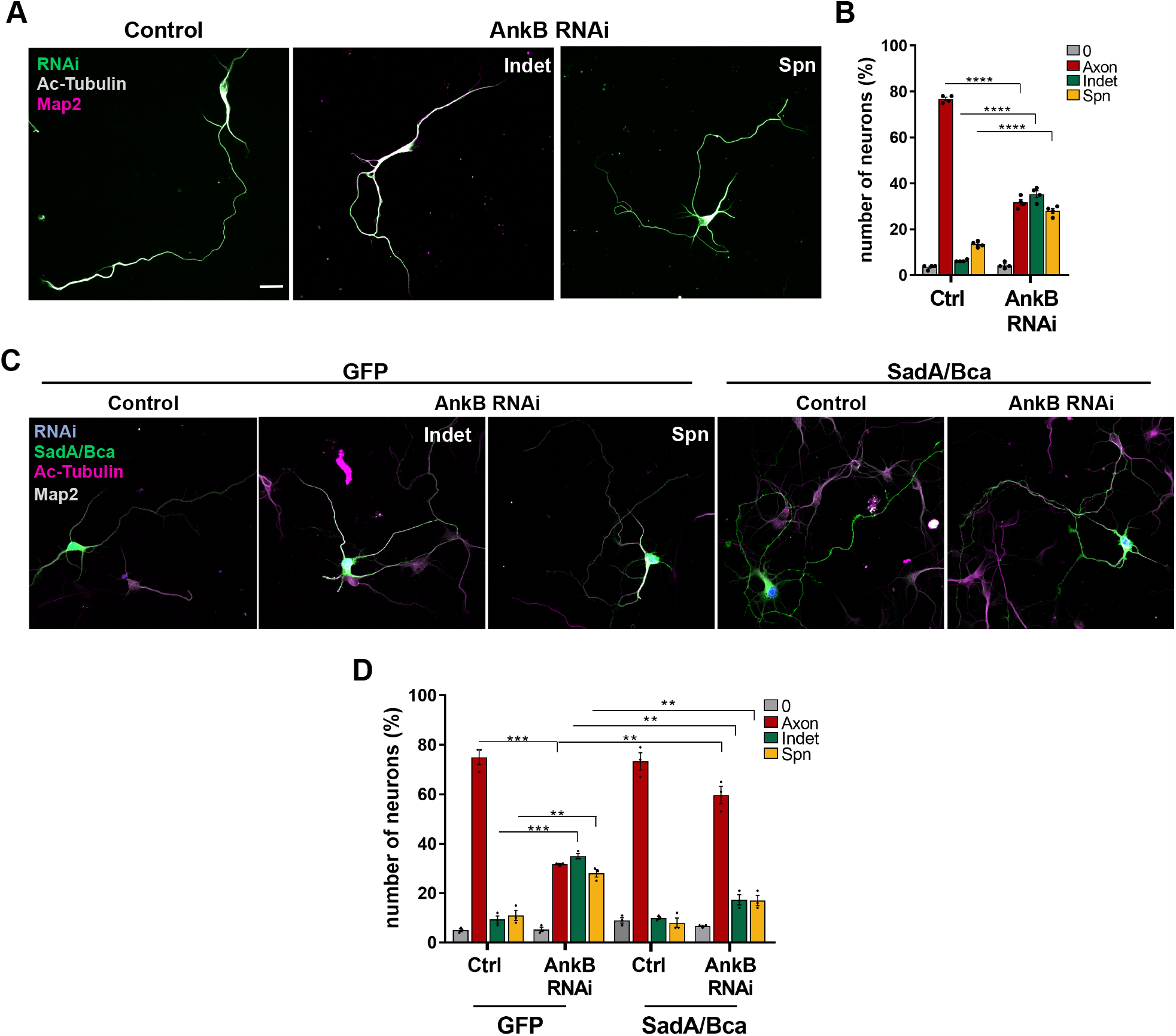
AnkB acts upstream of SadA/B. (A) Cortical neurons were transfected with control or AnkB knockdown vectors expressing GFP (AnkB RNAi, green) and stained with anti-acetylated tubulin (light gray) and -Map2 (red) antibodies. Scale bars: 20µm. (B) The percentage of unpolarized neurons without an axon (0), polarized neurons with a single axon (Axon), neurons with indeterminate neurites (Indet) and supernumerary axons (Spn) is shown for (A) (unpaired Student’s t-test; n=4 independent experiments). (C) Cortical neurons were transfected with control or AnkB knockdown vectors expressing H2B-mRFP (pseudo-colored blue) and GFP (control) or constitutively active SadA and SadB (SadA/Bca; green). Neurons were stained with anti-acetylated tubulin (magenta) and -Map2 (light gray) antibodies. Scale bars: 20µm. (D) The percentage of unpolarized neurons without an axon (0), polarized neurons with a single axon (Axon), neurons with indeterminate neurites (Indet) and supernumerary axons (Spn) is shown for (C) (unpaired Student’s t-test; n=3 independent experiments). Values are means ± SEM. See also Figure S2.

### The loss of SadA/B or AnkB increases the length of mitochondria

AnkB localizes to intracellular membranes including mitochondria and lysosomes, while phospho-T175/186-SadA/B is restricted to the axon but not to a specific organelle (Fig. S3A) (Dhumale et al., 2018; Galiano et al., 2012; Lorenzo et al., 2014; Qu et al., 2016). To determine if they are present on mitochondria we labeled them by expression of mitoGFP and stained neurons at 3 d.i.v. with antibodies for AnkB and phospho-T175/186-SadA/B. Signals for AnkB were detectable on 72±2% of the mitochondria in axons (Fig. S3B and S3C), confirming previous results (Qu et al., 2016). AnkB-positive sites were analyzed for the presence of phosphorylated SadA/B (Fig. 3A and 3B). 38±1% of the mitochondria showed at least one site positive for both phosphorylated SadA/B and AnkB. Moreover, we observed that 53±3% of mitochondria showed at least one site of colocalization with both AnkB and Lamp1, as marker for lysosomes (Fig. 3C and 3D) and 48±2% with both phosphorylated SadA/B and Lamp1 (Fig. 3E and 3F). This colocalization suggests that the interaction of AnkB with SadA/B might stimulate their kinase activity at sites of contact between mitochondria and lysosomes.

**Figure 3.**
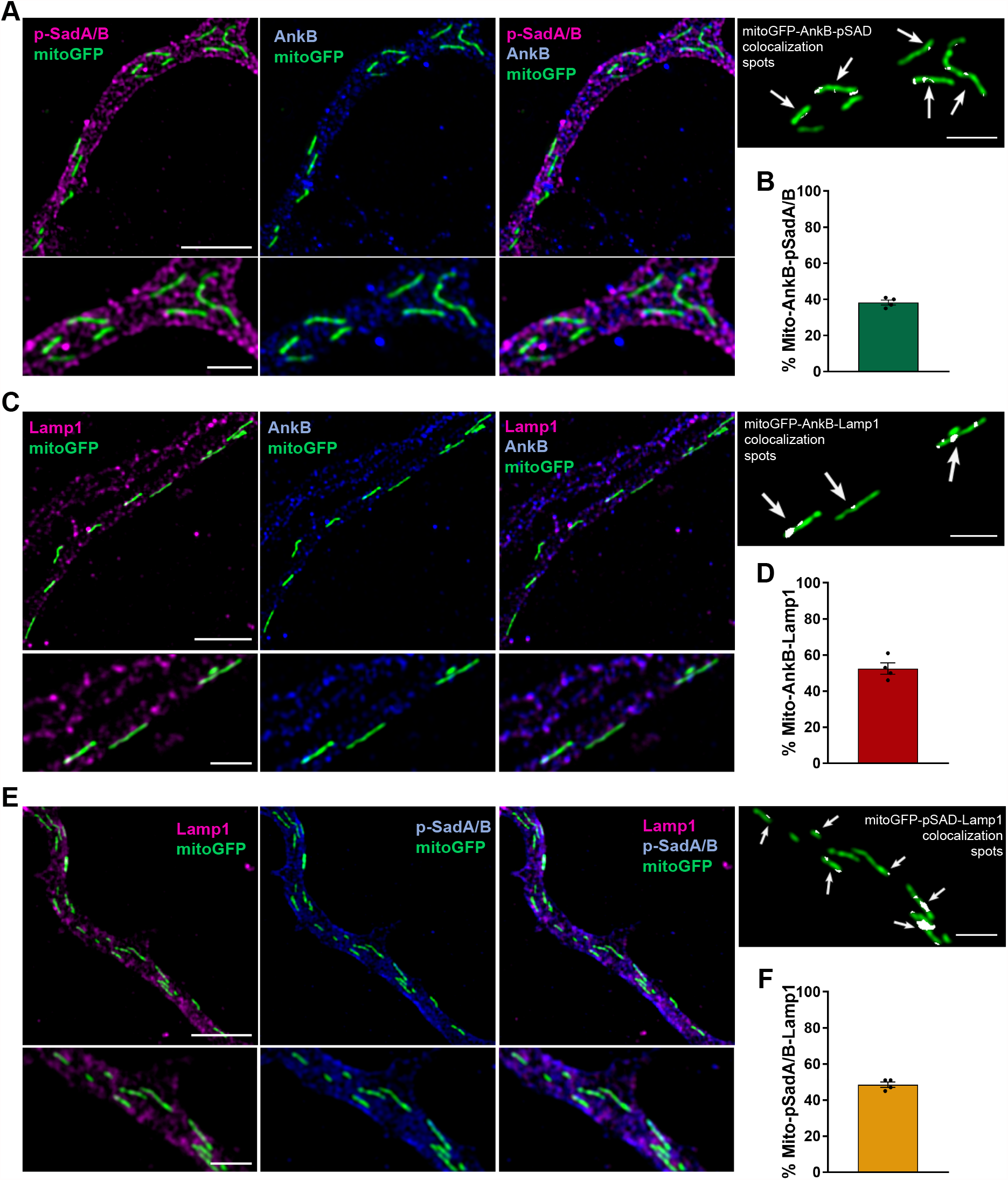
AnkB, phosphorylated SadA/B and Lamp1 colocalize at axonal mitochondria. (A) Neurons expressing mitoGFP (green) were stained with antibodies against phospho-T175/186-SadA/B (magenta) and AnkB (blue). (B) The percentage of mitochondria showing sites positive for both phospho-T175/186-SadA/B and AnkB is shown for (A). (C) Neurons expressing mitoGFP (green) were stained with antibodies against Lamp1 (magenta) and AnkB (blue). (D) The percentage of mitochondria showing sites positive for both Lamp1 and AnkB is shown for (C). (E) Neurons expressing mitoGFP (green) were stained with antibodies against Lamp1 (magenta) and phospho-T175/186-SadA/B (blue). (F) The percentage of mitochondria with sites positive for both LAMP1 and phospho-T175/186-SadA/B is shown for (E). Higher magnification Airyscan images are shown in the lower panels. White arrows indicate sites of colocalization at mitochondria. Scale bars: 5µm (upper panels), 2µm (lower panels) (N=90 neurons from n=4 independent experiments). Values are means ± SEM. See also Figure S3.

Contacts with lysosomes mark sites of mitochondrial fission (Wong et al., 2018, 2019; Yoshimori et al., 2018). Therefore, we analyzed the morphology of mitoGFP-labeled mitochondria at 3 d.i.v. after knockdown of both SadA and SadB. The average length of mitochondria in controls is 1.6±0.1µm in axons and 3.6±0.2µm in minor neurites (Fig. 4A, 4B and S4A). After knockdown of SadA and SadB, mitochondrial length was significantly increased to 3.2±0.2µm in indeterminate neurites and 3.0±0.1µm in supernumerary axons (Fig. 4A, 4B and S4B) that were analyzed separately. To exclude the possibility that the elongation of mitochondria after knockdown of SadA/B is a secondary consequence of the defect in the establishment of neuronal polarity, we transfected neurons at 1 d.i.v. instead of 0 d.i.v. (Fig. 4C). The majority of neurons (65±2% of the transfected H2B-mRFP-positive neurons) polarized normally and extended a single axon when transfected at this stage (Fig S4C). However, the length of axonal mitochondria was still significantly increased (Fig. 4D and S4D; control: 1.8±0.1, SadA/B RNAi: 2.7±0.1).

**Figure 4.**
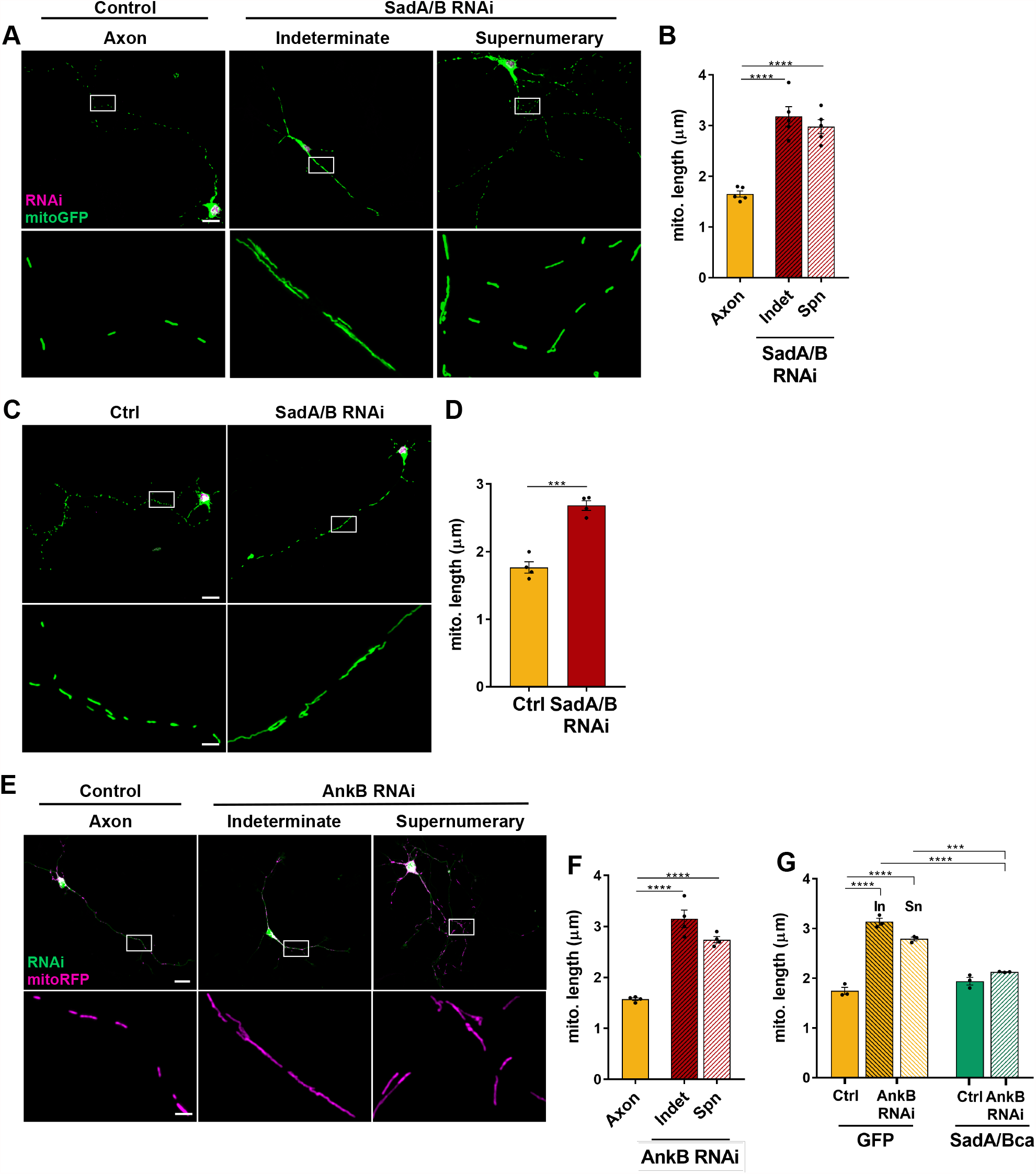
The loss of SadA/B or AnkB increases the length of mitochondria. (A) Cortical neurons were transfected with a vector for mitoGFP (green) and control or SadA/B knockdown vectors expressing H2B-mRFP (magenta). Higher magnification Airyscan images are shown in the lower panels. Scale bars: 20µm (upper panels), 2µm (lower panels). (B) The average mitochondrial length is shown for (A) (Indet: indeterminate neurites, Spn: supernumerary axons; one-way ANOVA followed by Dunnett’s test; N=125 neurons for each condition from n=5 independent experiments). (C) Cortical neurons were transfected at 1 d.i.v. and analyzed at 4 d.i.v. as in (A). Scale bars: 20µm (upper panels), 2µm (lower panels) (D) The average mitochondrial length is shown for (C) (unpaired Student’s t-test; N=100 neurons for each condition from n=4 independent experiments) (E) Cortical neurons were transfected with a vector for mitoRFP (magenta) and control or AnkB knockdown vectors expressing GFP (green). Scale bars: 20µm (upper panels), 2µm (lower panels). (F) The average mitochondrial length is shown for (E) (Indet: indeterminate neurites, Spn: supernumerary axons; one-way ANOVA followed by Dunnett’s test; N=100 neurons for each condition from n=4 independent experiments). (G) Quantification of average mitochondrial length in cortical neurons transfected with a vector for mitoRFP, control or AnkB knockdown vectors and GFP (control) or constitutively active SadA and SadB (SadA/Bca) (In: indeterminate neurites; Sn: supernumerary axon; one-way ANOVA followed by Tukey’s test; n=3 independent experiments). Values are means ± SEM. See also Figure S4.

After knockdown of AnkB, the length of axonal mitochondria was increased to a similar extent (Fig. 4E, 4F and S4E; control: 1.6±0.03µm, AnkB RNAi: 3.1±0.2µm (indeterminate neurites), 2.7±0.06 µm (supernumerary axons)). The co-expression of constitutively active SadA/B reversed the elongation and completely restored normal mitochondrial length after AnkB knockdown (Fig. 4G, S4F and S4G; AnkB RNAi: 3.1±0.07µm (indeterminate neurites), 2.8±0.04µm (supernumerary axons); AnkB RNAi +SadA/Bca: 2.1±0.01µm). Expression of SadA/Bca did not change the length of mitochondria in controls. Thus, Sad kinases are required for regulating the length of mitochondria downstream of AnkB independently of neuronal polarization.

### SadA/B are required to maintain the balance of mitochondrial fission and fusion

In cortical axons, the rates of mitochondrial fission and fusion are balanced to maintain their morphology (Cagalinec et al., 2013; Lewis et al., 2018; Saxton and Hollenbeck, 2012). An imbalance in these rates changes mitochondrial length. To determine if the elongation of axonal mitochondria after the knockdown of SadA/B or AnkB results from impaired mitochondrial dynamics, we performed live cell imaging of mitoGFP- or mitoRFP-labeled mitochondria in the proximal and middle segment of cortical axons at 3 d.i.v. (Suppl. Video S1 and S2). Our analysis revealed a striking increase in mitochondrial motility in SadA/B knockdown neurons. The percentage of motile mitochondria was increased in both the anterograde (Fig. 5A and 5E; control: 16±1%; SadA/B RNAi: 40±5% (indeterminate neurites), 28±4% (supernumerary axons)) and retrograde direction (control: 16±2%; SadA/B RNAi: 38±5% (indeterminate neurites), 33±4% (supernumerary axons)).

**Figure 5.**
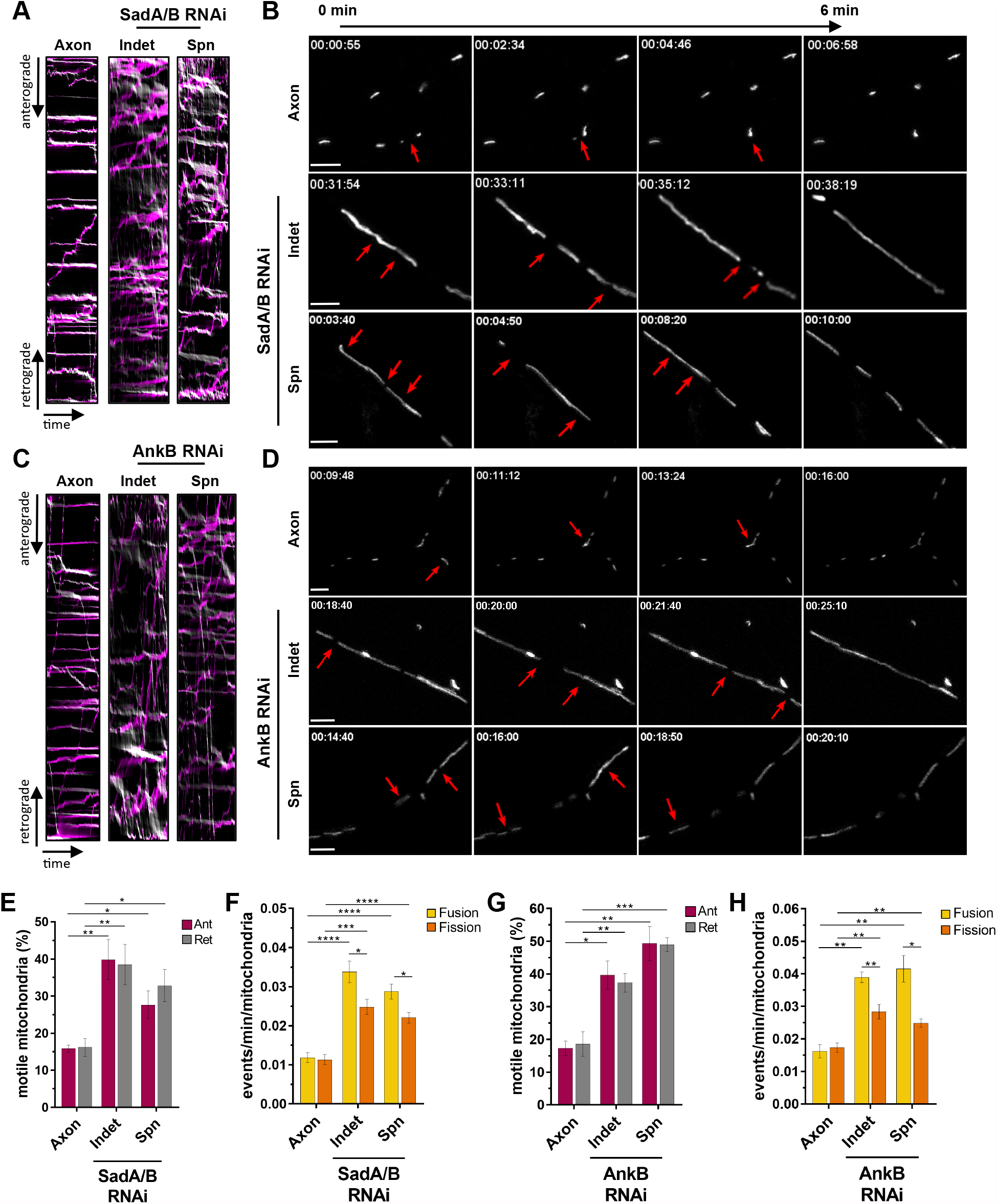
SadA/B are required to maintain the balance of fission and fusion. (A and C) Cortical neurons were transfected with a vector for mitoGFP (A and B) or mitoRFP (C and D) and control, SadA/B (A and B) or AnkB (C and D) knockdown vectors. Mitochondrial dynamics was analyzed by live cell imaging. Kymographs show anterograde (magenta) and retrograde (grey) transport of mitochondria in control (Axon) or knockdown neurons. (B and D) Selected frames from time-lapse videos show mitochondrial fission and fusion events from control (Axon), SadA/B (B) or AnkB (D) knockdown neurons. Red arrows mark fission or fusion events. Scale bars: 5µm. (E and G) The percentage of anterogradely (Ant) and retrogradely (Ret) moving axonal mitochondria after knockdown of SadA/B (E) or AnkB (G) is shown (N=22 (E) and 12 (G) neurons for each condition from n=6 (E) and n=3 (G) independent experiments) (F and H) Fission and fusion were quantified after knockdown of SadA/B (F) or AnkB (H) as the number of events/min/mitochondria (N=22 (F) and 12 (H) neurons for each condition from n=6 (F) and n=3 (H) independent experiments) (Indet: indeterminate neurites, Spn: supernumerary axons; one-way ANOVA followed by Dunnett’s test was done or all experiments). Values are means ± SEM. See also Figures S5 and Videos S1 and S2.

Next, we quantified the rate of mitochondrial fission and fusion as the number of events/min/mitochondria. In control neurons, the rate of fission (0.011±0.001) was similar to the rate of fusion (0.012±0.001). The knockdown of SadA/B resulted in a strong but unequal increase in the rates of both fission and fusion that disrupted their balance (Fig. 5B and 5F; indeterminate neurites: 0.034±0.003 (fusion), 0.025±0.002 (fission), supernumerary axons: 0.022±0.001 (fission), 0.029±0.002 (fusion)). This unequal increase leads to a reduction in the percentage of fission events as a fraction of all events (fission + fusion) compared to controls (Fig. S5A; control: 50±1% of all events; SadA/B RNAi: 41±1% (indeterminate neurites), 43±1% (supernumerary axons)).

Knockdown of AnkB resulted in a phenotype similar to that of SadA/B and induced an increase in both the number of motile mitochondria (Fig. 5C and 5G) and the rate of mitochondrial fission and fusion (Fig 5D and 5H; AnkB RNAi: 0.039±0.002 (fusion) and 0.030±0.002 (fission) in indeterminate neurites; 0.042±0.004 (fusion) and 0.028±0.001 (fission) in supernumerary axons). Similar to the SadA/B knockdown, the AnkB knockdown changed the balance of fission and fusion with a decrease in the percentage of fission events (Fig. S5B; control: 52±1%; AnkB RNAi: 42±2% (indeterminate neurites); 38±2% (supernumerary axons)). Taken together, these results show that the knockdown of SadA/B or AnkB, while increasing mitochondrial motility and dynamics, disturbs the balance of fission and fusion by decreasing the percentage of fission events, which results in an elongation of mitochondria.

### A decrease in Drp1 recruitment affects mitochondrial dynamics

Drp1 is a dynamin-related GTPase that oligomerizes on the OMM to induce fission (Cagalinec et al., 2013; Francy et al., 2015; Smirnova et al., 2001). Silencing of Drp1 results in a dramatic elongation of mitochondria. Therefore, we asked whether the relative reduction in mitochondrial fission after knockdown of SadA/B results from an impaired Drp1 recruitment to mitochondria. Staining with an anti-Drp1 antibody revealed Drp1 puncta both in the cytoplasm and on mitochondria at 3 d.i.v. (Fig. 6A) (Losón et al., 2013; Smirnova et al., 2001). The number of Drp1 puncta per mitochondria increases with their length in control neurons. While the average length of mitochondria is higher after knockdown of SadA/B, the number of Drp1 puncta per mitochondria was significantly reduced for every size class compared to controls (Fig. 6B). These results suggest that the knockdown of SadA/B impairs the recruitment of Drp1 to the OMM, which could explain the relative reduction in fission events.

**Figure 6.**
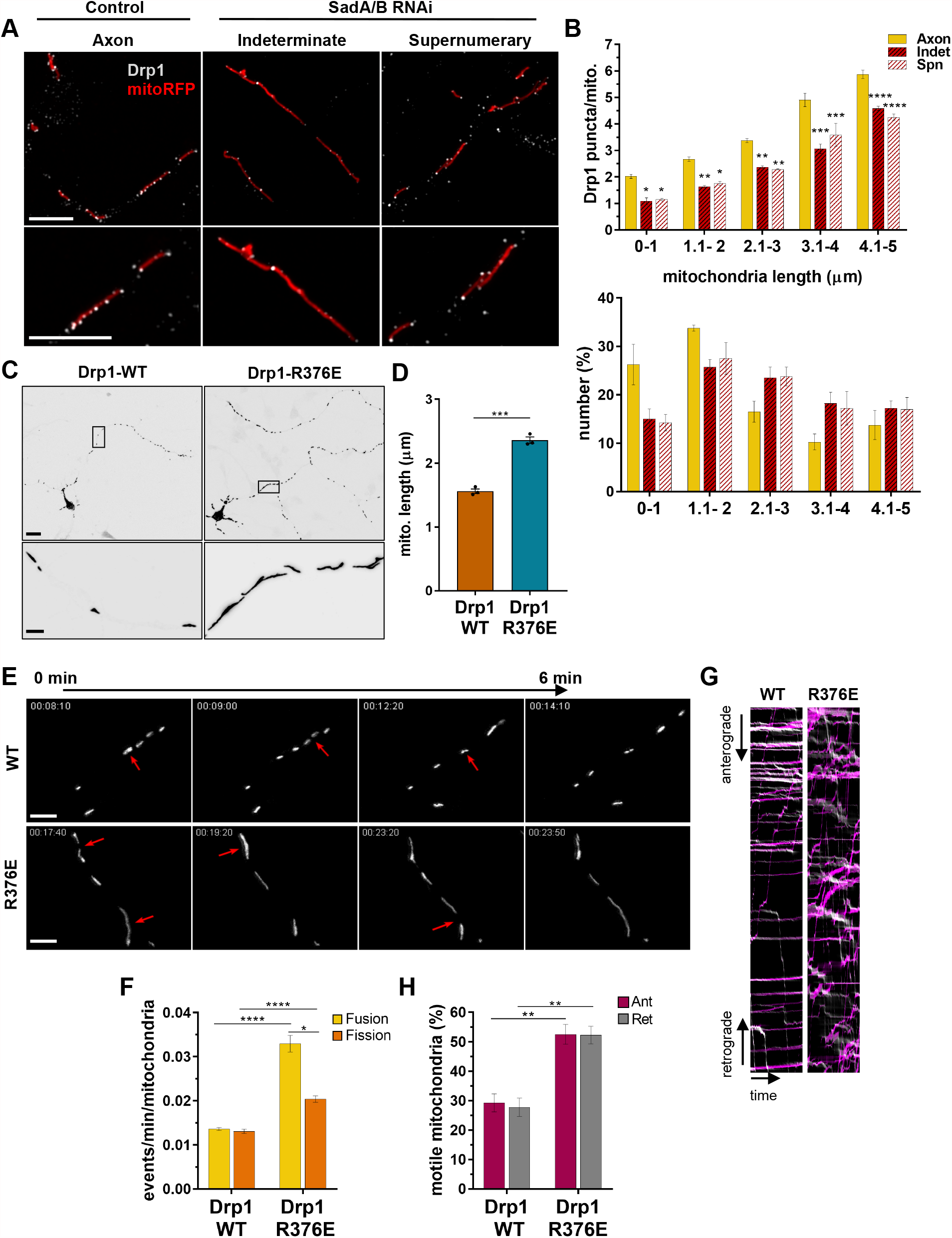
A decrease in Drp1 recruitment affects mitochondrial dynamics. (A) Cortical neurons were transfected with a vector for mitoRFP (red) and control or SadA/B knockdown vectors and stained with an anti-Drp1 antibody (white). Scale bars: 5µm. (B) The number of Drp1 puncta localizing to mitochondria was assessed by Manders’ colocalization coefficient-based analysis and is shown for different size classes of mitochondria (Indet: indeterminate neurites, Spn: supernumerary axons; one-way ANOVA followed by Tukey’s test; N=120 neurons for each condition for n=4 independent experiments). The total number of mitochondria for each size class is shown below. (C) Cortical neurons were transfected with vectors for mitoGFP and wild type Drp1 (Drp1 WT) or Drp1-R376E. Scale bars: 20µm (upper panels), 2µm (lower panels). (D) The average mitochondrial length is shown for (C) (unpaired Student’s t-test; N=75 neurons for each condition from n=3 independent experiments). (E) Selected frames from time-lapse imaging show mitochondrial fission and fusion events from cortical neurons transfected as in (C). Red arrows mark fission or fusion events. Scale bars: 5µm. (F) Fission and fusion events were quantified from (E) as the number of events/min/mitochondria (unpaired Student’s t-test; n=4 independent experiments). (G) Kymographs show anterograde (magenta) and retrograde (light gray) axonal transport of mitoGFP-labelled mitochondria in neurons expressing wild type Drp1 (WT) or Drp1-R376E. (H) The percentage of anterogradely (Ant) and retrogradely (Ret) moving axonal mitochondria is shown for (G) (unpaired Student’s t-test; n=4 independent experiments). Values are means ± SEM. See also Figures S6 and Video S3.

To determine whether impaired recruitment of Drp1 is sufficient to alter mitochondrial dynamics we employed Drp1 with a mutation in the stalk domain (Drp1-R376E) that reduces the interaction with its receptor Mff (Otera et al., 2010). Drp1-R376E blocks Drp1 association with mitochondria and impairs mitochondrial fission. The expression of Drp1-R376E did not affect neuronal polarity (Fig. S6A) but resulted in an elongation of axonal mitochondria (Fig. 6C, 6D and S6B; Drp1 WT: 1.5±0.03 µm, Drp1-R376E: 2.4±0.01 µm). Live cell imaging of Drp1-R376E-expressing neurons (Suppl. Video S3) showed a higher number of both mitochondrial fission and fusion events with a lower increase in fission than fusion (Fig. 6E and 6F; 0.0329±0.002 (fission); 0.0204±0.001 (fusion)) compared to wild type Drp1 (0.0130±0.0005 (fission); 0.0136±0.0003 (fusion)). This unequal change results in a lower percentage of fission events compared to wild type Drp1 (Fig. S6C; Drp1 WT: 49±0.8%; Drp1-R376E 38±1.3%). The percentage of motile mitochondria was also increased in both the antero- and retrograde direction by Drp1-R376E expression (Fig. 6G and 6H). In summary, the expression of Drp1-R376E induced a change in mitochondrial dynamics that is very similar to that observed after the knockdown of SadA/B kinases.

### Tau regulates mitochondrial length downstream of SadA/B

Overexpression of Tau results in defects in mitochondrial morphology due to an inhibition of Drp1 localization to mitochondria (DuBoff et al., 2012). In order to examine if SadA/B regulate mitochondrial size through the phosphorylation of Tau, we tested if expression of phosphomimetic Tau-S262D, Tau-S356D or Tau-S262/356D is able to rescue the defects in SadA/B knockdown neurons. Expression of phosphomimetic but not wild type Tau was able to restore neuronal morphology in SadA/B knockdown neurons, as predicted by previous results (Barnes et al., 2007; Kishi et al., 2005; Rodríguez-Asiain et al., 2011; Yoshida and Goedert, 2012) while it had no effect in controls (Fig. 7A; indeterminate neurites: 43±1% (SadA/B RNAi), 21±1% (RNAi+Tau-S262D), 21±1% (RNAi+Tau-S356D), 12±1% (RNAi+Tau-S262/356D); supernumerary axons: 33±2% (SadA/B RNAi), 15±1% (RNAi+Tau-S262D), 17±1% (RNAi+Tau-S356D), 13±1% (RNAi+Tau-S262/356D). Notably, also mitochondrial length was fully normalized in SadA/B knockdown neurons by phosphomimetic but not wild type Tau (Fig. 7B, 7C, S7A and S7B; SadA/B RNAi 3.2±0.1 µm (indeterminate neurites), 2.8±0.2 µm (supernumerary axons); RNAi+Tau-S262D: 1.6±0.04 µm; RNAi+Tau-S356D: 1.5±0.1 µm; RNAi+Tau-S262/356D: 1.5±0.1 µm). Tau constructs mimicking phosphorylation of a single Serine residue were already sufficient for a rescue. The restoration of mitochondrial length suggests that SadA/B kinases regulate the size of mitochondria by phosphorylating Tau.

**Figure 7.**
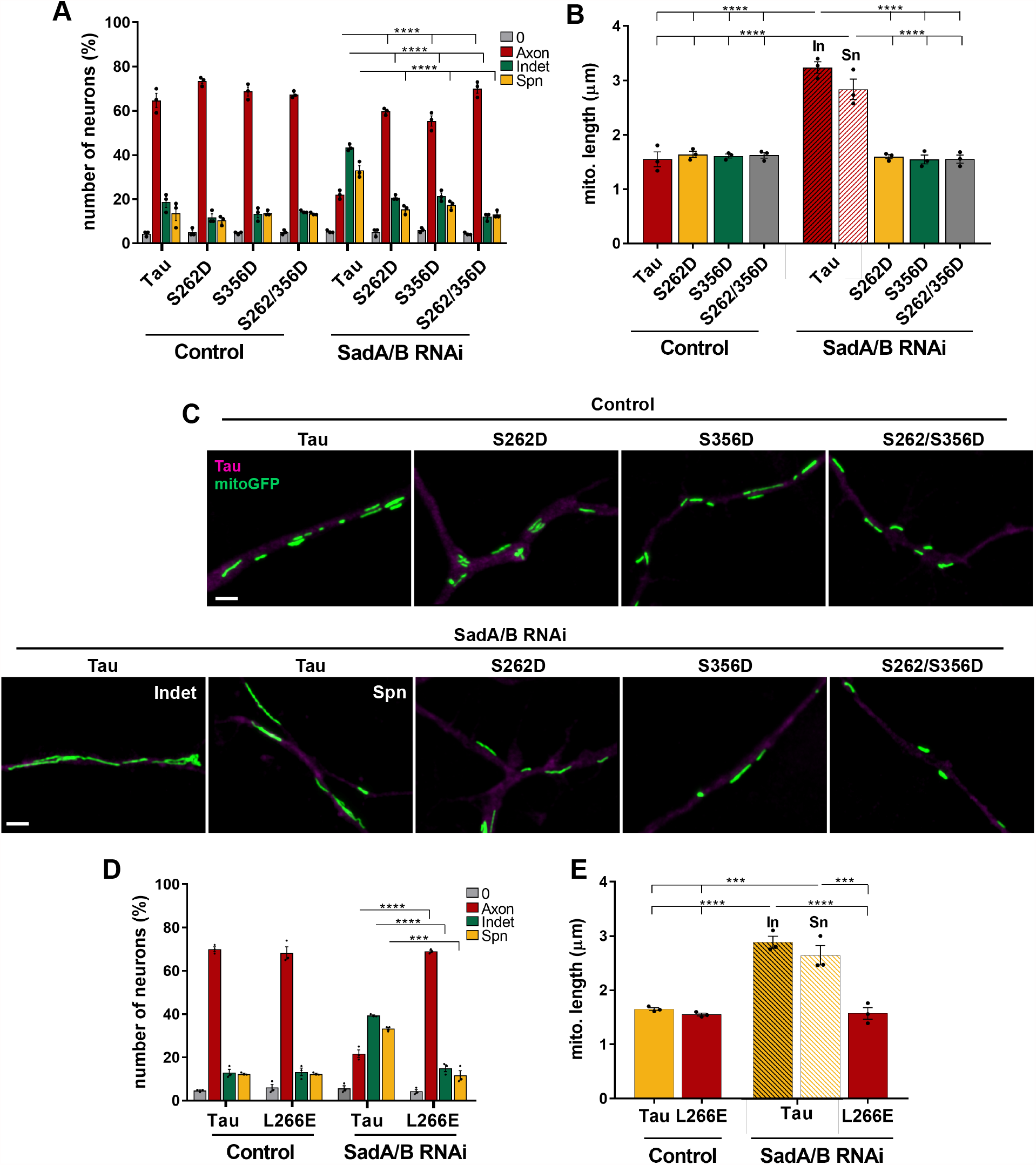
Tau regulates mitochondrial length downstream of SadA/B. (A) Cortical neurons were transfected with control or SadA/B knockdown vectors and vectors for wild type or phosphomimetic Tau as indicated. The percentage of unpolarized neurons without an axon (0), polarized neurons with a single axon (Axon), neurons with indeterminate neurites (Indet) and supernumerary axons (Spn) is shown (one-way ANOVA followed by Tukey’s test; n=3 independent experiments). (B) The average mitochondrial length is shown for (A) (Indet: indeterminate neurites, Spn: supernumerary axons; one-way ANOVA followed by Dunnett’s test; n=3 independent experiments) (C) Higher magnification Airyscan images of mitoGFP (green) and wild type or phosphomimetic Tau (magenta) are shown for (B). Scale bars: 2µm (D) Cortical neurons were transfected with control or SadA/B knockdown vectors and a vector for wild type Tau or Tau-L266E. The percentage of unpolarized neurons without an axon (0), polarized neurons with a single axon (Axon), neurons with indeterminate neurites (Indet) and supernumerary axons (Spn) is shown (unpaired Student’s t-test; n=3 independent experiments). (E) The average mitochondrial length is shown for (D) (In: indeterminate neurites, Sn: supernumerary axons; one-way ANOVA followed by Tukey’s test; n=3 independent experiments). Values are means ± SEM. See also Figures S7.

It was previously shown that Tau interacts with and stabilizes F-actin (Cabrales Fontela et al., 2017; Elie et al., 2015; Fulga et al., 2007; He et al., 2009). Both excessive stabilization and complete depolymerization of actin filaments inhibits Drp1 translocation to mitochondria, resulting in mitochondrial elongation (DuBoff et al., 2012; Ji et al., 2015; Korobova et al., 2013, 2014). The phosphorylation of Ser262 and Ser356 in the repeat domain of Tau reduces its affinity for F-actin (Cabrales Fontela et al., 2017). Thus, decreased Tau phosphorylation might affect mitochondrial dynamics due to increased binding and excessive stabilization of actin filaments. To address this possibility, we tested if expression of Tau-L266E with a reduced affinity for F-actin can also rescue the knockdown of SadA/B (Cabrales Fontela et al., 2017). Expression of Tau-L266E restored neuronal polarity but did not affect its establishment in control neurons (Fig. 7D; indeterminate neurites: 40.3±1% (SadA/B RNAi), 15±1% (RNAi+Tau-L266E); supernumerary axons: 33±1% (SadA/B RNAi), 12±2% (RNAi+Tau-L266E)). Expression of Tau-L266E in SadA/B knockdown neurons significantly reduced mitochondrial length in neurons with a single axon to a value similar to controls (Fig. 7E, S7C and S7D; SadA/B RNAi: 2.9±0.1 µm (indeterminate neurites), 2.6±0.2 µm (supernumerary axons); SadA/B RNAi+Tau-L266E: 1.6±0.1 µm). Thus, the elongation of mitochondria upon knockdown of SadA/B can be rescued by expression of a Tau mutants that binds F-actin with decreased affinity.

### Actin destabilization rescues mitochondrial defects after knockdown of SadA/B

To demonstrate that mitochondrial defects in SadA/B knockdown neurons arise from excessive F-actin stability, we attempted to restore normal mitochondrial length by mildly destabilizing actin filaments with low doses of Latrunculin A (LatA). LatA sequesters actin monomers but also destabilizes actin filaments by severing and promoting depolymerization (Fujiwara et al., 2018). Cortical neurons were transfected with SadA/B knockdown vectors and treated with 1 nM LatA. Unlike higher doses of LatA (Dupraz et al., 2019), low doses did not affect the establishment of polarity in control neurons. LatA also did not reduce the number of neurons with indeterminate neurites or supernumerary axons after knockdown of SadA/B (Fig. 8A). However, the elongation of mitochondria after loss of SadA/B could be fully reversed by LatA treatment for 17h at 2 d.i.v. or for 5h at 3 d.i.v. (Fig 8B, S8A, S8B and S8C; indeterminate neurites: 3±0.1 µm (SadA/B RNAi), 2±0.1 µm (SadA/B RNAi+17h LatA), 2.1±0.1 µm (SadA/B RNAi+5h LatA); supernumerary axons: 2.9±0.2 µm (SadA/B RNAi), 1.8±0.1 µm (SadA/B RNAi+17h LatA), 1.8±0.1 µm (SadA/B RNAi+5h LatA)).

**Figure 8.**
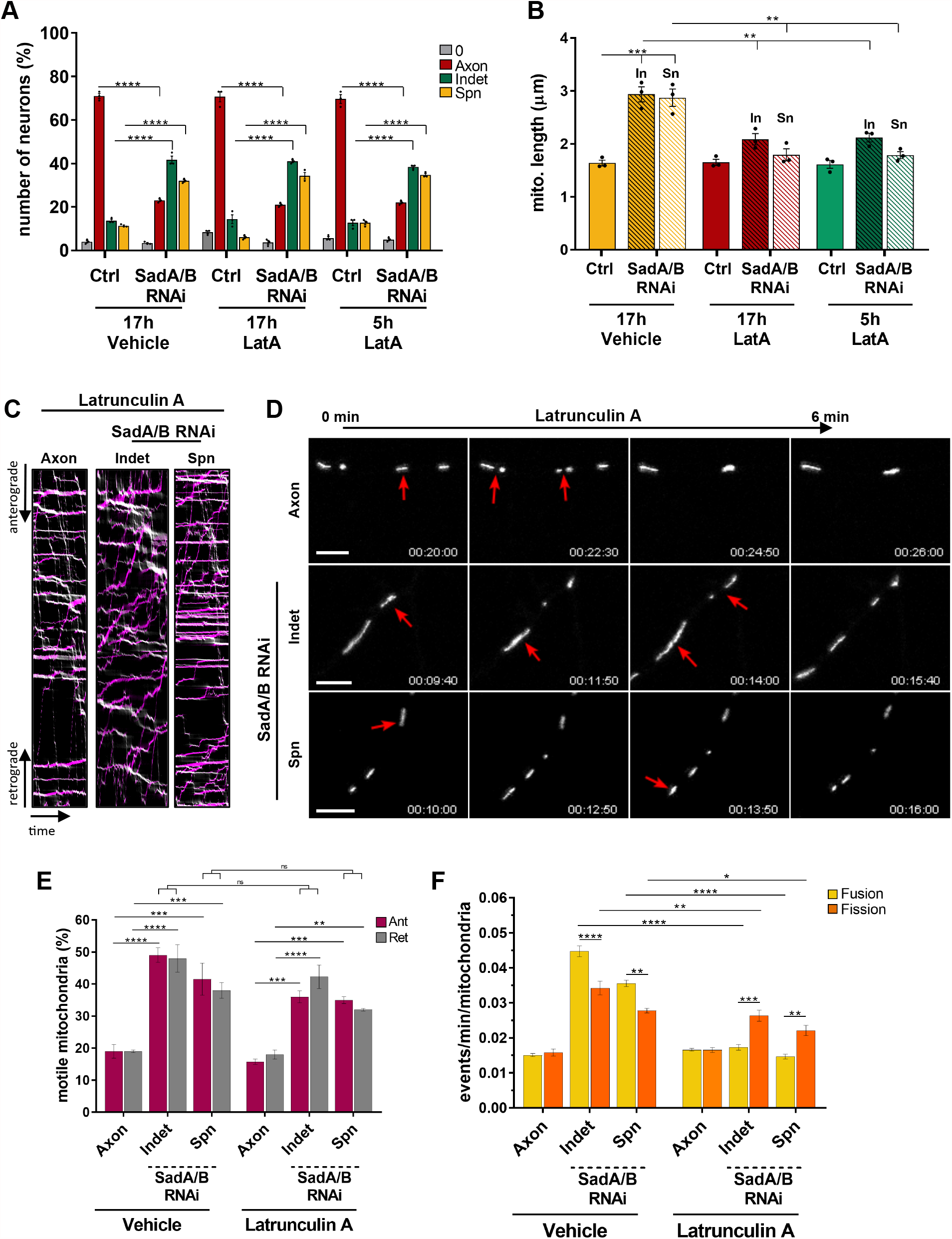
Actin destabilization rescues mitochondrial defects after SadA/B knockdown. (A) Cortical neurons were transfected with control or SadA/B knockdown vectors and treated with vehicle (DMSO) for 17h or 1nM LatA for 17h and 5h as indicated. The percentage of unpolarized neurons without an axon (0), polarized neurons with a single axon (Axon), neurons with indeterminate neurites (Indet) and supernumerary axons (Spn) is shown (one-way ANOVA followed by Tukey’s test; n=3 independent experiments). (B) The average mitochondrial length is shown for (A) (In: indeterminate neurites. Sn: supernumerary axons; one-way ANOVA followed by Dunnett’s test; n=3 independent experiments). (C) Cortical neurons were transfected with control or SadA/B knockdown vectors and treated with vehicle (DMSO) or 1nM LatA for 5h. Mitochondrial dynamics was analyzed by live cell imaging. Kymographs show anterograde (magenta) and retrograde (light gray) axonal transport of mitochondria in control (Axon) or SadA/B knockdown neurons treated with 1nM LatA for 5h. (D) Selected frames from time-lapse imaging show mitochondrial fission and fusion events from control (Axon) and SadA/B knockdown neurons treated with 1nM LatA for 5h (Indet: indeterminate neurites; Spn: supernumerary axons). Red arrows mark fission or fusion events. Scale bars: 5µm. (E) The percentage of anterogradely (Ant) and retrogradely (Ret) moving axonal mitochondria are shown for (C) (one-way ANOVA followed by Tukey’s test; n=4 independent experiments). (F) Fission and fusion events were quantified from (D) as the number of events/min/mitochondria (Indet: indeterminate neurites, Spn: supernumerary axons; one-way ANOVA followed by Tukey’s test; n=4 independent experiments). Values are means ± SEM. See also Figures S8 and Video S4.

To confirm that the reduction in mitochondrial length after actin destabilization is due to changes in the rates of fission and fusion, we analyzed mitochondrial motility and dynamics in neurons treated at 3 d.i.v. for 5 hours with 1 nM LatA (Suppl. Video S4). The percentage of motile mitochondria was not significantly lower than in controls with vehicle after this short treatment (Fig. 8C and 8E). LatA does not alter the balance between mitochondrial fission and fusion in control neurons. However, the number of fission and fusion events was significantly reduced in SadA/B knockdown neurons treated with LatA (Fig. 8D and 8F; SadA/B RNAi: 0.045±0.002 (fusion) and 0.034±0.002 (fission) in indeterminate neurites; 0.028±0.0007 (fission) and 0.036±0.0009 (fusion) in supernumerary axons); SadA/B RNAi+LatA: 0.026±0.001 (fission) and 0.017±0.0008 (fusion) in indeterminate neurites; 0.022±0.001 (fission) and 0.015±0.0007 (fusion) in supernumerary axons). LatA changed the balance of fission and fusion and increased the percentage of fission events compared to controls (Fig. S8D; control: 50±1%; indeterminate neurites: 62±1%; supernumerary axons: 60±1%). These results strongly suggest that the defects in mitochondrial dynamics after suppression of SadA/B are due to excessive actin stabilization.

## Discussion

As dynamic organelles, mitochondria continuously change their morphology and distribution while maintaining a relatively uniform length in axons through balanced rates of fission and fusion (van der Bliek et al., 2013; Cagalinec et al., 2013; Lewis et al., 2018; Seager et al., 2020; Tilokani et al., 2018). In this study we show that the kinases SadA and SadB function downstream of AnkB to regulate mitochondrial dynamics in cortical axons. We observed a striking increase in mitochondrial length after the loss of SadA/B due to alteration in the balance of fission and fusion. These changes result from excessive actin stabilization and decreased Drp1 localization to mitochondria due to reduced Tau phosphorylation. SadA/B are, therefore, required for the characteristic size of axonal mitochondria in cortical neurons. They join AMPK, Nuak1 and Mark2 that regulate various aspects of mitochondrial homeostasis (Courchet et al., 2013, 2018; Matenia et al., 2012; Toyama et al., 2016).

SadA and SadB are required for the differentiation of distinct axonal and dendritic extensions during neuronal development (Barnes et al., 2007; Dhumale et al., 2018; Kishi et al., 2005; Müller et al., 2010). The elongation of mitochondria after their loss could, therefore, arise as a secondary consequence of defects in axon formation. However, changes in mitochondrial dynamics were detected not only in indeterminate neurites but also in supernumerary axons formed after SadA/B knockdown. Moreover, the same mitochondrial elongation can be observed in normally polarized neurons when SadA/B were suppressed after axons are specified. Finally, mild actin destabilization rescued the imbalance in the rate of fission and fusion and normalized the length of mitochondria but did not restore neuronal polarity. Taken together, these results indicate that mitochondrial dynamics is directly affected by the loss of SadA/B independently of an effect on other axon-specific properties.

Live cell imaging revealed that the suppression of SadA/B disturbs the balance between fission and fusion by a shift towards a lower percentage of fission events. While both, fission and fusion events, were more frequent, this increase was unequal resulting in an elongation of mitochondria. The loss of SadA/B reduced the localization of Drp1 to mitochondria, which can explain the lower rate of fission. The recruitment and oligomerization of Drp1 to mitochondria is a critical step for fission and Drp1 deletion results in a striking elongation of mitochondria in axons (Berthet et al., 2014). Expression of the Drp1-R376E mutant with decreased affinity for the mitochondrial receptor Mff results in a similar phenotype as the knockdown of SadA/B. Thus, impairing the recruitment of Drp1 is sufficient to alter mitochondrial dynamics and length without affecting the polarization process. Fission and fusion of mitochondria do not occur independently of each other and are coordinated at contacts sites with the ER where the machinery for both processes is present (Abrisch et al., 2020; Guo et al., 2018). The reduced recruitment of Drp1 may, therefore, have consequences also for the fusion of mitochondria.

It has been established before that Tau is a substrate for SadA/B (Barnes et al., 2007; Kishi et al., 2005; Rodríguez-Asiain et al., 2011; Yoshida and Goedert, 2012). Our experiments confirmed that Tau phosphorylation at Ser356 is reduced after their knockdown. Expression of phosphomimetic Tau was able to restore both neuronal polarity and normal mitochondrial length in SadA/B knockdown neurons, confirming that Tau is a major target for SadA/B in the developing nervous system required for both functions. Actin filaments promote Drp1 oligomerization and are essential for mitochondrial fission (Fenton et al., 2021; Moore, 2018). Tau binding was shown to excessively stabilize actin filaments in a model for tauopathy leading to impaired Drp1 localization (DuBoff et al., 2012). Tau phosphorylation at Ser262 and Ser356 attenuates the interaction with both microtubules and F-actin (Cabrales Fontela et al., 2017; Fulga et al., 2007). Therefore, a deficit in Tau phosphorylation could increase its binding and stabilization of actin filaments.

Depolymerization of actin by short-term treatment with high doses of Latrunculin B reduces mitochondrial fission and increases the length of mitochondria (Korobova et al., 2013, 2014). However, treatment of SadA/B knockdown neurons with low doses of LatA to mildly destabilize actin filaments has the opposite effect and restores the relatively small and uniform size of mitochondria by increasing the percentage of fission events. These experiments provide direct evidence for the conclusion that the increase in mitochondrial length after knockdown of SadA/B arises from excessive F-actin stabilization. Actin filaments cycle through the mitochondrial network in HeLa cells by transiently assembling on subpopulations of mitochondria (Moore et al., 2016). This cycling of actin would be impaired by excessive stabilization by Tau since it requires a dynamic actin network, but it remains to be shown that it can be observed also in axons.

In addition to the elongation of mitochondria, a striking increase in the motility of mitochondria was observed after the knockdown of SadA/B that affected both antero- and retrograde transport. LatA treatment did not significantly alter mitochondrial motility in control or SadA/B knockdown neurons. The increased motility may, therefore, require higher LatA doses for a rescue or depend on an actin-independent mechanism. The transport of mitochondria along axonal microtubules is driven by kinesin-1 and dynein (Fenton et al., 2021). Tau inhibits the transport by kinesin-1 by steric hindrance but has little effect on dynein (Chaudhary et al., 2018; Dixit et al., 2008; Ebneth et al., 2011; Monroy et al., 2018, 2020; Siahaan et al., 2019; Tan et al., 2019; Vershinin et al., 2007). However, an increased binding of Tau to microtubules due to a loss of SadA/B would be expected to decrease mitochondrial motility rather than increase it. Only a fraction of mitochondria is usually motile in axons, while the majority is stationary and anchored to microtubules by factors like Syntaphilin (Franker and Hoogenraad, 2013; Kang et al., 2008; Pathak et al., 2010). Syntaphilin inactivation results in a dramatic increase in their motility (Kang et al., 2008). Since the binding mode of Syntaphilin to the microtubule lattice remains to be characterized, it cannot be excluded that an increased localization of Tau to microtubules reduces the proportion of anchored mitochondria. The increase in the rate of motility and fusion suggests that SadA/B regulate several parameters of mitochondrial dynamics through additional mechanisms that remain to be elucidated in future studies.

We identified AnkB as an upstream regulator of SadA/B that interacts with and stimulates their kinase activity as revealed by the phosphorylation of Tau. Knockdown of AnkB results in similar defects in neuronal morphology and mitochondrial size as a SadA/B knockdown and can be rescued by constitutively active SadA/B. AnkB associates with mitochondria and lysosomes in neurons (Lorenzo et al., 2014; Qu et al., 2016). Contact sites between those organelles mark the position of mitochondrial fission events (Wong et al., 2018, 2019; Yoshimori et al., 2018). A high percentage of mitochondria showed signals for AnkB that colocalized with phosphorylated SadA/B and both colocalized also with a lysosomal marker. The activation of SadA/B is, therefore, restricted to a limited number of sites along mitochondria, which may correspond to contacts between mitochondria and lysosomes based on their colocalization. Since phosphorylated SadA/B is not restricted to mitochondria, additional interaction partners are likely to regulate SadA/B activation at other sites (Hung et al., 2007; Ma et al., 2016; Wu et al., 2015; Yang et al., 2012). Our findings suggest that AnkB regulates mitochondrial motility and dynamics by promoting the activation of SadA/B in addition to its function as an adaptor for the dynein-dynactin motor complex (Lorenzo et al., 2014).

Taken together, our results suggest that AnkB acts upstream of SadA/B in regulating mitochondrial motility and the balance of fission and fusion. SadA/B phosphorylate Tau and modulate mitochondrial dynamics by regulating the actin-dependent recruitment of Drp1 to mitochondria. These results identify a pathway in which SadA/B promote the acquisition of distinct properties of axonal mitochondria.

## Acknowledgments

We thank Maria Wenning and Ina Kowsky for technical assistance and M. Bähler for plasmids. This work was supported by the Deutsche Forschungsgemeinschaft (DFG) through a grant to A.W.P. (PU-102-12), the Cells-in-Motion Cluster of Excellence (EXC 1003-CiM) and SFB 1348.

## Author Contributions

A.P. conceived the study, D.D.M., P.R. and P.D. performed the experiments, D.D.M., P.R., P.D. and A.P. contributed to the experimental design and interpretation of the results, and D.D.M., P.R. and A.P. wrote the manuscript.

## STAR Methods

### LEAD CONTACT AND MATERIALS AVAILABILITY

Further information and requests for resources and reagents should be directed to and will be fulfilled by the corresponding author, Dr. Andreas Püschel. All unique/stable reagents generated in this study are available from the corresponding author without restriction.

### EXPERIMENTAL MODEL AND SUBJECT DETAILS

All animal protocols were performed in accordance with the guidelines of the North Rhine-Westphalia State Environment Agency (Landesamt für Natur, Umwelt und Verbraucherschutz (LANUV)). Rats were maintained at the animal facility of the Institute for Molecular Cell Biology (University of Münster) under standard housing conditions at four to five per cage with a 12 h light/dark cycle (lights on from 07:00 to 19:00 h) at constant temperature (23°C) with continuous access to food and water. Timed-pregnant rats were set up in house. Pregnant rats were anesthetized by exposure to isoflurane followed by decapitation and primary cultures prepared from E18 embryos

## METHOD DETAILS

### Neuronal Cell Culture and Transfection

Primary cortical neurons were obtained from embryonic day 18 (E 18) rat embryos and transfected by calcium phosphate co-precipitation. In brief, cortices were dissected, collected in HBSS (Invitrogen) and digested in trypsin (0.25% Trypsin/EDTA, Invitrogen) for 8 min. Digested tissue was mechanically dissociated in DMEM (1X DMEM (Invitrogen), 2mM glutamine, 50U/mL gentamycin, 10% fetal calf serum (FCS)) and passed through a cell strainer (40 µm pore size, Sarstedt). Dissociated neurons were seeded onto poly-L-ornithine coated (10 µg/ml, SigmaAldrich) coverslips at a density of 35,000 cells/cm^2^ for immunofluorescence staining on poly-L-ornithine coated 35mm dish (µ-Dish Quad, Ibidi) at a density of 150,000 cells/cm^2^ for live cell imaging. Neurons were allowed to attach in the presence of DMEM for 2 h. The medium was then replaced by Neurobasal medium ((1x NBM (Invitrogen), 2mM glutamine, 50U/mL gentamycin, 2% B27 (v/v)) or BrightCell^TM^ NEUMO photostable medium (1X NEUMO (Merck Millipore), 4% SOS, 4 mg/ml human recombinant insulin, 2mM glutamine, 50U/mL gentamycin) for staining or live cell imaging, respectively. For transfection, the culture medium was replaced 3 h after plating by Opti-MEM (Invitrogen) before the DNA mixture was added. After incubation for 45 min at 37°C at 5% CO_2_, neurons were washed for 15 min with 1 ml of Opti-MEM, which had been pre-incubated at 37°C at 10% CO_2_, and the conditioned Neurobasal medium was added back to the cells. Cortical neurons were transfected at 0 d.i.v. and fixed at 3 d.i.v. if not indicated otherwise.

For analysis by Western blot, primary cortical neurons were transfected by nucleofection using *Nucleofector*™ *II* (Amaxa) and the Ingenio® Electroporation kit (Mirus Bio LLC). For one well of a 6-well plate, 1.5 × 10^6^ cells were resuspended in 100 µL Ingenio® electroporation solution with 3 µg of plasmid DNA and nucleofection was performed using the program O-003. For analysis of phospho-S356-Tau levels, neurons were lysed in RIPA buffer 72 h after seeding.

### Plasmids

The following plasmids were obtained from Addgene: pAnkyrinB-2HA (#31057); pLV-RFP (#26001); pmRuby2-MAPTau-C-10 (#55904); pmCh-Drp1 (#49152); psfGFP-C1 (#54579). Red FP Vector-Mitochondrion (mitoRFP (#558722)) was purchased from BD Pharmingen. Mitochondrion-EGFP (mitoGFP) was a gift from Martin Bähler (Oeding et al., 2018).

The expression vectors for EGFP-SadA-WT, EGFP-SadB-WT, EGFP-SadA-KI (S179D), EGFP-SadB-KI (S191D), EGFP-SadAca (T175E) and EGFP-SadBca (T187E) have been described before (Müller et al., 2010).

The constructs expressing the Kinase-UBA (SadA: 19-342 aa; SadB: 33-358 aa) or AIS-KA1 (SadA: 343-636 aa; SadB: 359-715 aa) (Fig. 1A) domains of SadA/B were prepared from full-length mouse SadA and SadB using the primers described in Table S1. The PCR products were cloned into the psfGFP-C1.

Loss of function experiments were performed with miRNAs generated using the BLOCK-iT Pol II miRNA Expression Vector Kit (Invitrogen) (Table S1). Target sequences were cloned into the pcDNA6.2-GW/EmGFP-miR expression plasmid following the manufacturer’s instructions. To generate pcDNA6.2-GW/H2B-mRFP-miR, H2B-mRFP was excised from pLV-RFP with DraI/SalI and inserted into pcDNA6.2-GW/EmGFP to replace the EmGFP sequence.

Single point mutations were generated by site directed mutagenesis using Phusion High-Fidelity DNA Polymerase (Thermo Fisher) (Table S2). Tau mutants (Tau-S262D, Tau-356D, Tau-262/356D, Tau-S262A, Tau-S356A, Tau-S262/356A and Tau-L266E) were generated from pmRuby2-MAPTau-C-10. The Drp1-R376E mutant was generated from pmCh-Drp1. Non-phosphorylatable SadA and SadB mutants (SadA-T175A, SadB-T186A) were generated from pEGFP-SadA and pEGFP-SadB. All the primers were purchased from Metabion.

### Immunofluorescence staining

Neuronal cultures were fixed with 4% paraformaldehyde and 4% sucrose for 15 min at 37°C and quenched in 50 mM ammonium chloride at room temperature (RT) for 10 min. After several washes with Phosphate-Buffered Saline (PBS), the cells were permeabilized with 0.1% Triton X-100 for 3 min and treated with blocking solution (2% normal goat serum, 2% bovine serum albumin and 0.2% fish gelatine in PBS) for 1 h at RT. Fixed cells were incubated overnight with the primary antibodies (listed in the key resources table) at 4°C. Cells were washed with PBS and incubated with the appropriate Alexa-Fluor conjugated secondary antibody (Thermo Fisher) at RT for 2 h and mounted using Mowiol (SigmaAldrich). Antibodies were diluted in 10% blocking solution.

### Transfection of HEK 293T cells and Western blot

HEK293T cells were transfected using the calcium phosphate co-precipitation method. Transfected HEK293T cells were harvested in ice-cold PBS and lysed at 4°C for 1 h (Lysis buffer: Tris/HCl 50 mM, pH 7,4, NaCl 150 mM, DTT 1 mM, MgCl2 1,5 mM, EDTA 4 mM, glycerol 10% (v/v), Triton X-100 1% (v/v), complete protease inhibitor (Roche), tyrosine and serine/threonine phosphatase inhibitors (Sigma). After addition of 5x Laemmli buffer and boiling at 95°C, proteins were separated by SDS-polyacrylamide gel electrophoresis and transferred to nitrocellulose membranes. Non-specific binding was blocked at RT with 5% non-fat dry milk or 3% fish gelatin (for detection of phosphorylated proteins, SigmaAldrich) in Tris-Buffered Saline and 0.1% Tween20 (TBS-T) for 1 h. The membrane was incubated overnight with primary antibodies diluted in blocking solution at 4°C. After several washes with TBS-T, membranes were incubated with HRP-coupled secondary antibodies (Jackson ImmunoResearch Labs) for 2h at RT. Peroxidase activity was visualized by the enhanced chemiluminescence detection system (Interchim) using the ChemiDocTM MP imaging system (Bio-Rad).

### Immunoprecipitation with anti-GFP nanobodies

For co-immunoprecipitation experiments, a GST-tagged anti-GFP nanobody was expressed in E. coli BL21 cells as described (Katoh et al., 2015). The anti-GFP nanobody was coupled to Glutathione Sepharose™ 4B beads (GE Healthcare) and incubated with lysates from transfected HEK 293T cells for 1 h at 4°C. The beads were washed three times (10 minutes each with Tris/HCl pH 7.4 50 mM, NaCl 250 mM, DTT 1 mM, MgCl2 1.5 mM, EDTA 4 mM, glycerol 10% (v/v), Triton X-100 0.1% (v/v), complete protease inhibitor). Bound proteins were eluted using 2x Laemmli buffer and analyzed by western blotting.

### Image acquisition and analysis

Images were acquired on a Zeiss LSM 800 Airyscan laser scanning confocal microscope with 405, 488, 568, and 647 nm laser lines and processed using the Zeiss ZEN 2.3 (blue edition) software (Carl Zeiss MicroImaging, Jena, Germany). Neuronal and mitochondrial morphology was analyzed in fixed neurons using a Plan-Apochromat 40x/1,3 Oil DIC M27 objective. Co-localization studies were performed with a Plan-Apochromat 40x/1,3 Oil DIC M27 objective using images acquired with the Airyscan detector (Carl Zeiss MicroImaging, Jena, Germany). Time-lapse imaging for mitochondrial dynamics analysis was performed in a CO_2_ regulated incubation chamber maintained at 37°C (PeCon, Erbach). Images were acquired at a frame rate of one scan every 10 seconds for 25-45 minutes. Images were analyzed using ImageJ (NIH).

## QUANTIFICATION AND STATISTICAL ANALYSIS

### Quantification

Neuronal polarity was analyzed at 3 d.i.v as described previously (Dhumale et al., 2018). Neurons showing acetylated-tubulin-negative neurites that were positive for MAP2 were categorized as unpolarized and neurons with a single long and acetylated-tubulin-positive neurite as normally polarized (Witte et al., 2008). Neurons with several long and acetylated-tubulin-positive neurites showing accumulation of acetylated-tubulin were counted as neurons with supernumerary axons and those with several neurites of comparable length and positive for both, axonal (acetylated-tubulin) and dendritic markers (Map2), as neurons with indeterminate neurites. At least 150 cells per condition were counted for each experiment.

Mitochondrial length was quantified in selected ROIs (scanning zoom: 1.2-1.5X) with resolvable individual mitochondria. User-defined thresholds for pixel intensity were used on maximum intensity projections of mitoGFP or mitoRFP stacks to identify measurable mitochondria and make binary images. The length of single mitochondria was then obtained by using the analyze particles tool. The analysis was performed on equal numbers of neurons for each condition in all experiments. The number of mitochondria moving along the axon was quantified from color-coded kymographs that were generated using the macro toolset KymographClear in ImageJ (Mangeol et al., 2016). The rate of fusion and fission was quantified by frame-by-frame manual analysis. For the analysis of colocalization, Airyscan images were pre-processed by filtering and adjusting the fluorescence threshold to eliminate diffuse signal. The same settings were used for all images. Colocalization spots were identified using the JaCoP plug-in (Bolte and Cordelières, 2006) for ImageJ (Manders’ coefficient) and then manually counted.

Colocalization analysis for three channels was done on Airyscan images. Each channel was first processed by filtering and background subtraction to reduce noise. The same settings were used for all images of each channel. Images were segmented by thresholding in order to get binary masks that were finally combined using the image calculator tool (operation: AND) in order to obtain triple-colocalization masks. The latter were used to identify the number of mitochondria that show one site of colocalization with AnkB and phospho-Thr175/Thr186-SadA/B or Lamp1, respectively.

### Statistical Analysis

All plotted values are expressed as means ± standard error of the mean (SEM). All experiments were performed independently at least three times. Specific numbers can be found in the figure legends. If not otherwise indicated, comparisons between two groups were performed using unpaired Students t-test, comparisons of more than two groups were done using one-way ANOVA followed by Tukey’s or Dunnett’s test with GraphPad Prism. For all analyses performed, significance was defined as ns: p>0.05; *p≤0.05; **p≤0.01; ***p≤0.001; ****p≤0.0001. When not indicated, the comparison between conditions is not significantly different.

## Supplemental videos

**Video S1**. Knockdown of SadA/B increases the mitochondrial motility and disturbs the balance of fission and fusion. Related to Figure 5A and 5B.

**Video S2**. Knockdown of AnkB increases the mitochondrial motility and disturbs the balance of fission and fusion similar to SadA/B knockdown. Related to Figure 5C and 5D.

**Video S3**. Expression of Drp1-R376E increases the mitochondrial motility and disturbs the balance of fission and fusion in single axon neurons, similar to SadA/B and AnkB knockdown. Related to Figure 6E and 6G.

**Video S4**. LatA treatment of SadA/B knockdown neurons reduces the rate of fission and fusion and increases the percentage of fission events but does not restore mitochondrial motility. Related to Figure 8C and 8D.

